# Powerful Inference with the D-statistic on Low-Coverage Whole-Genome Data

**DOI:** 10.1101/127852

**Authors:** Samuele Soraggi, Carsten Wiuf, Anders Albrechtsen

## Abstract

The detection of ancient gene flow between human populations is an important issue in population genetics. A common tool for detecting ancient admixture events is the D-statistic. The D-statistic is based on the hypothesis of a genetic relationship that involves four populations, whose correctness is assessed by evaluating specific coincidences of alleles between the groups.

When working with high throughput sequencing data calling genotypes accurately is not always possible, therefore the D-statistic currently samples a single base from the reads of one individual per population. This implies ignoring much of the information in the data, an issue especially striking in the case of ancient genomes.

We provide a significant improvement to overcome the problems of the D-statistic by considering all reads from multiple individuals in each population. We also apply type-specific error correction to combat the problems of sequencing errors and show a way to correct for introgression from an external population that is not part of the supposed genetic relationship, and how this leads to an estimate of the admixture rate.

We prove that the D-statistic is approximated by a standard normal. Furthermore we show that our method outperforms the traditional D-statistic in detecting admixtures. The power gain is most pronounced for low/medium sequencing depth (1-10X) and performances are as good as with perfectly called genotypes at a sequencing depth of 2X. We show the reliability of error correction on scenarios with simulated errors and ancient data, and correct for introgression in known scenarios to estimate the admixture rates.

## Introduction

An important part in the understanding of a population’s history and its genetic variability is past contacts with other populations. Such contacts could result in gene flow and admixture between populations and leave traces of a population’s history in genomic data. In fact, the study of gene flow between populations has been the basis to uncover demographic histories of many species, including human and archaic human populations [2-5,8,12-15,22,23,33].

The study of the history of human populations using both modern and ancient human genomes has become increasingly topical with the recent availability of new high-throughput sequencing technologies [6], such as Next Generation Sequencing (NGS) technologies [7]. These technologies have made it possible to obtain massive quantities of sequenced DNA data even from ancient individuals, such as an Anzick-Clovis individual from the Late Pleistocene [8], a Neandertal individual [2] and a Paleoamerican individual [9].

There are many different methods for inferring and analyzing admixture events using genome-scale data. Popular methods such as `STRUCTURE` [10] and `ADMIXTURE` [11] estimate how much a sampled individual belongs to *K* clusters that often can be interpreted as the individual’s admixture proportion to the *K* populations. However, these approaches are not appropriate to detect ancient gene flow and do not work well with a limited number of individuals per population.

A recent alternative to the above methods is the D-statistic. The D-statistic is based on the di-allelic patterns of alleles between four groups of individuals, and provides a way to test the correctness of a hypothetical genetic relationship between the four groups (see Fig 1). A variant of the D-statistic (called the *F*_4_-statistic) was first used in [12] to identify that subgroups of the Indian Cline group are related to external populations in term of gene flow. Also the amount of gene flow might be estimated using the *F*_4_-statistic [4].

**Fig 1.**
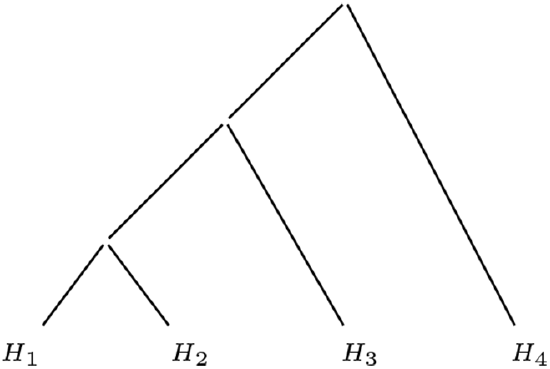
Tree topology for the D-statistic. Hypothesis of genetic relationship between four populations *H*_1_,*H*_2_,*H*_3_,*H*_4_.

In the pivotal study [2] the D-statistic was used to show that 3 non-African individuals are more genetically similar to the Neandertal sequence than African San and Yoruban individuals are. Moreover, it has been shown that the Eastern Asian populations have a higher amount of Neandertal shared genetic material [4].

Using the D-statistic on many Old World and Native Americans it has been suggested gene flow into some Native American populations, such as evidence of admixture from Australasian populations into New World Populations [22,33].

In another study the affinity between the Anzick genome and the Native Americans genome was analyzed with the D-statistic to compare different hypotheses regarding their ancestry [8]. Using the D-statistic, it has been reported that the remains of an individual from the Mal’ta population in south-central Siberia have contributed to the gene pool of modern-day Native Americans, with no close affinity to east Asians [13].

The first use of the D-statistic was based on a sampling approach that allowed to perform the test without the need to call SNPs or genotypes [2]. This approach is still widely used, and amongst the available computational tools implementing this approach is the `doAbbababa` program of `ANGSD` [16] (supporting low depth NGS data) or the `fourpop` program of `TreeMix` [17] (supporting di-allelic genotype data and microsatellite data). The program `qpDstat` of `ADMIXTOOLS` [15] computes the D-statistic from populations with multiple individuals from di-allelic genotype data. The program `doAbbababa` relies on sampling one base from every locus, using the sequenced reads to define the sampling probabilities.

The D-statistic is often applied to scenarios involving ancient individuals, that are commonly affected by deamination, i.e. the natural degradation of DNA after death of the organism that leads to there being few molecules remaining in ancient specimens and often results in a low sequencing depth. Furthermore, deamination can cause high frequency of specific transitions of the bases, low quality of the SNPs and very low depth of the data. The current methods for the D-statistic can be very ineffective and unreliable when applied to ancient data, since both sampling and genotype calling procedures are subject to high uncertainty.

The focus of this paper is to address the problems stated above. We propose a D-statistic - implemented in the program `doAbbababa2` of `ANGSD` - that supports low depth NGS data and is calculated using all reads of the genomes, and therefore allows for the use of more than one individual per group. We prove that the improved D-statistic is approximated by a standard normal distribution, and using both simulated and real data we show how this approach greatly increases the sensitivity of gene-flow detection and thus improves the reliability of the method, in comparison to sampling a single read. We also illustrate that it is possible to correct for type-specific error rates in the data, so that the reads used to calculate the D-statistic will not bias the result due to type-specific errors. Moreover, our improved D-statistic can remove the effect of known introgression from an external population into either *H_1_, *H*_2_* or *H_3_*, and indirectly estimates the admixture rate.

## Materials and Methods

This section introduces the traditional D-statistic and the theory that leads to its approximation as a normal distribution. Thereafter we explain how to extend the D-statistic to use multiple individuals per population, without genotype calling and still preserving the same approximation property of the D-statistic. Lastly, we will show how to deal with type-specific errors and introgression from a population external to the tree topology.

## Standard D-statistic

The objective of the D-statistic is to assess whether the tree of Fig 1 that relates four present-day populations *H*_1_,*H*_2_,*H*_3_,*H*_4_, is correct. When *H*_4_ is an outgroup, the correctness of the tree corresponds to the absence of gene-flow between *H*_3_ and either *H*_2_ or *H*_1_. This objective is achieved by developing a statistical test based on the allele frequencies and a null hypothesis *H*_0_ saying that the tree is correct and without gene flow. We limit the explanation to a di-allelic model with alleles A and B to keep the notation uncluttered; the extension to a 4-allelic model is fairly straightforward. We do not make assumption on which allele is derived, but we assume that B is the non-outgroup allele. Population *H*_4_ is an outgroup, that splits off at the root of the tree from the other branches. For each population *H_j_*, *j* = 1,2, 3,4, in the tree, we consider the related allele frequencies *x_j_*.

For each population *H_j_*, the observed data consists of a certain number of individuals sequenced without error. At every locus *i* there are 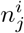 sequenced bases observed from aligned reads. We consider only the *M* loci for which there is at least one sequenced base from aligned reads in all four groups. Moreover, in this theoretical treatment we allow the number *M* of loci to grow to infinity. Assume that at a locus *i* the allele frequencies in the four groups of individuals 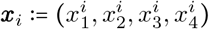 and let 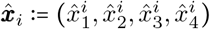 be an unbiased estimator of ***x***_*i*_, such as the relative frequencies of the allele A in each population.

The D-statistic focuses on di-allelic sites where the differences are observed within the pairs (*H*_1_, *H*_2_) and (*H*_3_, *H*_4_). Consider a random allele drawn from each of the four groups of genomes and the resulting combination of the four alleles. We are interested in two patterns:

- ABBA, meaning that we have the same allele in populations *H*_1_ and *H*_4_ and another allele from the individuals in populations *H*_2_ and *H*_3_;
- BABA, where an allele is shared by individuals in populations *H*_1_ and *H*_3_ and the other allele by individuals in populations *H*_2_ and *H*_4_.

The tree of Fig 1 is subject to independent genetic drifts of the allele frequencies along each of its branches. Consequently the probabilities of ABBA and BABA patterns conditionally to population frequencies would rarely be same. Therefore it is interesting to focus on their expected values with respect to the frequency distribution:

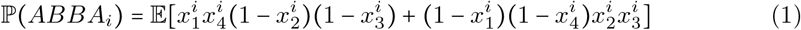

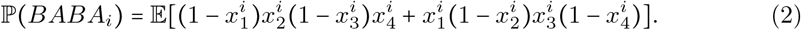

To verify that allele A is shared between genomes in *H*_1_, *H*_3_ as often as it happens between genomes in *H*_2_, *H*_3_, we require as null hypothesis that at each *i*-th locus the probability (1) equals the probability (2). This condition can be written as

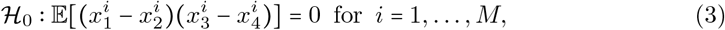

where the expectation is the difference between eq 1 and eq 2.

Using the empirical frequencies of the alleles as unbiased estimators for the population frequencies, we define the D-statistic as the following normalized test statistic

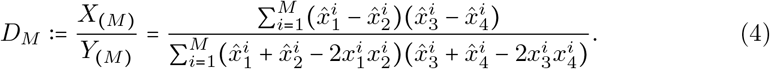

The values *X*_(*M*)_ and *Y*_(*M*)_ are the numerator and denominator, respectively. Using *Y*_(*M*)_ to normalize the numerator leads to the interpretation of *D_M_* as difference over all loci of the probabilities of having an ABBA or a BABA events, conditional to the event that only ABBA or BABA events are possible.

Appendix 1 shows that, under the hypothesis *H*_0_, the test statistic can be approximated by a standard normal variable. Specifically, the approximation holds with a proper rescaling, since *D_M_* would narrow the peak of the Gaussian around zero for large *M* (note that this rescaling is an embedded factor in the estimation of the variance of *D_M_* using the block jackknife method [21] in the software implementation of ANGSD). More generally the treatment could be extended to blockwise independence of the allele counts to take into account linkage disequilibrium.

The convergence results of Appendix 1 apply to the following special cases of the D-statistic:

1. the original D-statistic *D_M_* calculated by sampling a single base from the available reads [2] to estimate the sampling probabilities,
2. the D-statistic *D_M_* evaluated by substituting the frequencies 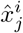 with the estimated population frequencies 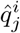 defined in eq 5 for multiple individuals (see Appendix 2).
3. the D-statistic *D_M_* evaluated only over loci where the outgroup is mono-allelic, such as when the Chimpanzee is set as an outgroup to test for gene flow from the Neandertal population into modern out-of-Africa populations [2].

## Multiple individuals per group

The D-statistic defined in eq 4 is calculated using population frequencies. In case only one individual per population is chosen, it is easy to get an estimator of the populations’ frequencies by simply counting observed bases. In what follows we are interested in getting a meaningful estimate of the frequencies in the case we want to use all the available sequenced individuals without calling genotypes.

This is done using a weighted sum of the estimated allele frequencies for each individual in every group. Assume that given the allele frequency 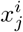, *j* = 1, 2,3,4, at locus *i* for the *j*th population, we model the observed data as independent binomial trials with parameters 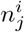 and 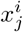, where 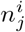 is the number of trials. We take the frequency of allele A in the reads of each *j*th population as an unbiased estimator of the population frequency. Let *N_j_* be the number of individuals in population *j*. For the *ℓ*th individual within the *j*th population, let 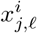 be the frequency of allele A at locus *i*, with estimator 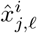 the frequency of allele A for ℓ = 1,…, *N_j_*. Define 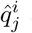 as the weighted sum

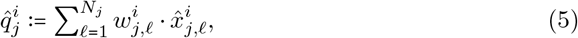

where each 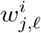 is a weight, that is proportional to a quantity depending on 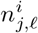, the number of sequenced bases at locus *i* for individual *ℓ*:

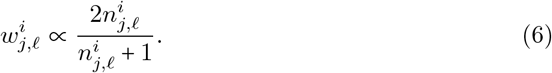

The estimator 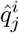 in eq (5) is an estimator for the *j*th population frequency at locus *i* with minimal variance (the derivation of the weights as minimizer of the frequency estimator’s variance can be found in Appendix 2). Substituting the estimated population frequencies in eq (4) with the weighted estimators determined by eq (5), it is possible to account for multiple individuals per population. Since the weighted estimator is also unbiased, it does not affect the approximation of the D-statistic with a standard normal distribution.

A first application of this method has been the estimation of population frequencies to reveal signatures of natural selection [18]. The weights have a strong impact on loci with low number of reads, where they assume a low value, leading to a stronger impact of population frequency estimated from high-depth individuals in each group.

## Error estimation and correction

The study of genetic relationships between populations often involves the use of ancient genomes that are subject to high error-rates. We introduce error correction following the idea illustrated in [19] to take errors into account and to obtain a more reliable D-statistic.

The estimation of the type specific error rates is possible using two individuals (one affected by type-specific errors, and one sequenced without errors) and an outgroup, denoted by T, R and O, respectively. Those individuals are considered in the tree ((T,R),O) (see Appendix 3).

After the error matrix is estimated for each individual it is possible to obtain error-adjusted frequencies of alleles in locus *i* through the following matrix-vector product:

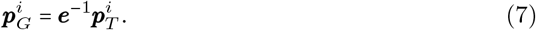

where 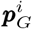 and 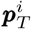 are the true and observed vectors of allele frequencies locus *i*, respectively, and **e**(*a, b*) - the type-specific error matrix with entries *a, b* in the set of possible alleles - is considered to be invertible. Note that estimating **e** and correcting the allele frequencies is a process best applied before the calculation of weighted allele frequencies for multiple individuals.

Using error-corrected estimators of the population frequencies to calculate the D-statistic does not prevent it to be approximated by a standard normal, because the error-corrected estimators are unbiased for the true population frequency (see Appendix 3).

According to eq (7) one is able to perform the error correction at every locus for every individual. In this way it is possible to build a weighted frequency estimator for each population after the error correction. However the implementation of eq (7) involves the inversion of a matrix and a matrix-vector multiplication at every locus for each individual in all populations. Moreover, as a consequence of the error estimation, there might be negative entries of the inverse **e**^-1^, which might cause the product of eq (7) to result in negative entries in the vector 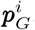.

Consequently we have decided to implement a less precise version of the error correction that is applied to each whole group of individuals instead of every single individual. Assume that the populations’ frequencies have been estimated from eq (5), and that it is possible to estimate the probabilities of the 256 alleles combinations AAAA, AAAC,…, TTTT between the four populations.

In each *j*th population of individuals, let **e**_(*j*)_ be the mean of their error matrices. Then build the error matrix for the four groups, ***E***. This has dimension 256 x 256 and its entry (*a*_1:4_,*b*_1:4_), where *a*_1:4_ = (*a*_1_,*a*_2_,*a*_3_,*a*_4_) and *b*_1:4_ = (*b*_1_, *b*_2_, *b*_3_, *b*_4_) are two possible allele patterns of the four populations, is defined as the probability of observing *b*_1:4_ instead of *a*_1:4_, assuming independence of the error rates between the four populations:

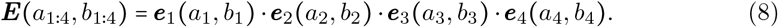

The equation states that the change from pattern *a*_1:4_ to *b*_1:4_ happens with a probability that is the product of the error rates of each population. Note that each error rate is the sum of the error rates of each individual in that population, and so does not take into account how every individual is weighted according to the frequency estimator of eq (5).

Let ***P***_*error*_ be the vector of length 256 that contains the estimated probabilities of observing allele patterns between the four populations, affected by type-specific errors. Denote by ***P***_*corr*_ the vector containing the estimated probabilities of patterns not affected by errors. With an approach similar to the one leading to eq 7 it holds that

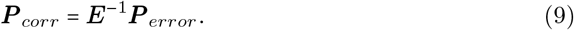

Using the error-corrected estimated probabilities of combinations of alleles of the type ABBA and BABA it is then possible to calculate numerator and denominator of the D-statistic. This procedure is fast but has the drawback that in every group the error matrix takes into account every individual within a population without its associated weight of eq 6. This means that the portion of alleles related to individuals with lower weights might undergo an excessive error correction.

## Correction for introgression from an external population

The improved D-statistic proves to be very sensitive to introgression, but a hypothesized genetic relationship might be rejected because of an admixture involving a population not part of the considered tree. We propose a way to correct this issue and obtain an estimate of the amount of introgression when the source of gene-flow is available.

In this section we analyze the case in which the null hypothesis might be rejected in favour of the alternative hypothesis, but the cause of rejection is not the presence of gene flow between *H*_3_ and either *H*_1_ or *H*_2_, but instead gene flow between an external population *H*_5_ and either *H*_2_ or *H*_1_. Consider the case of Figure S6A, where the null hypothesis might be rejected because of introgression from an external population *H*_5_ into *H*_2_ with rate a. We assume that the external sample for *H*_5_ represents the population that is the source of introgression. Consider *H*_2_ being the population subject to introgression from *H*_5_, and define 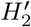 the same population when it has not undergone admixture.

The four population subtrees of interest (see Supplementary Figure 8) are *T*_1:4_ = *H*_2_)*H*_3_)*H*_4_), which includes the 4-population tree excluding the admixing population, *T_out_* = *H*_5_)*H*_3_)*H*_4_), where the population source of introgression replaces the admixed population, and 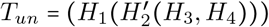, in which *H*_2_ has not yet undergone admixture and therefore reflects the null hypothesis *H*_0_.

**Figure 8.**
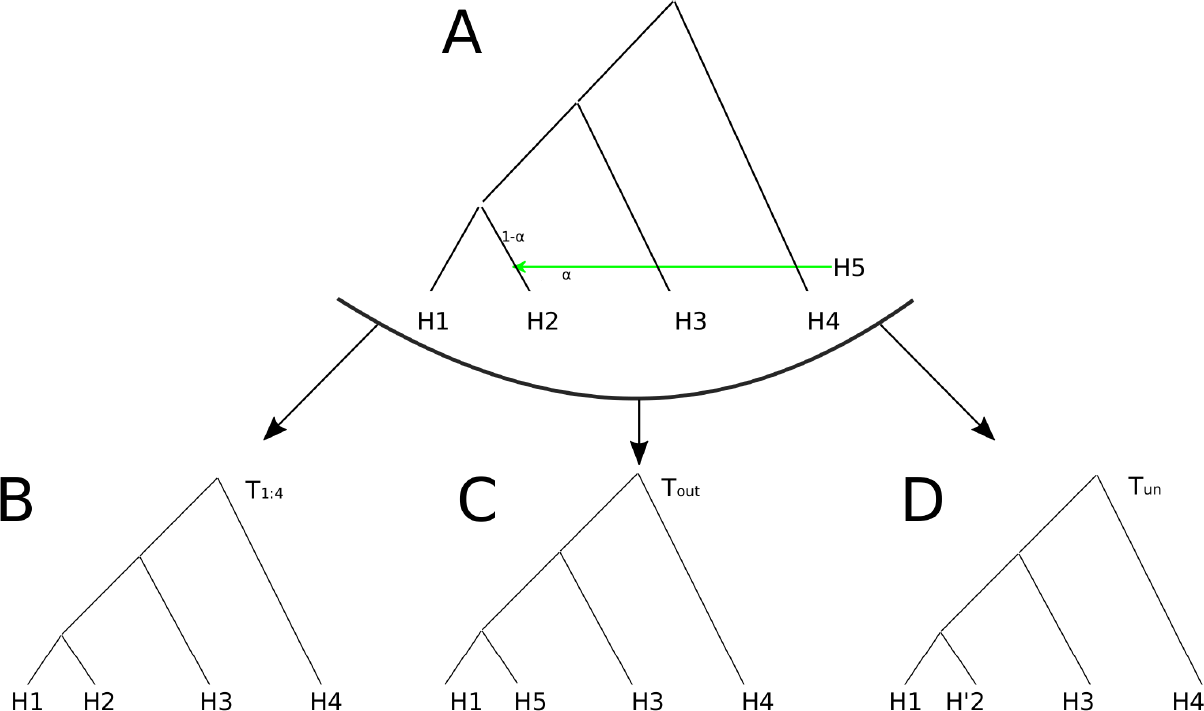
Subtrees of interest in a scenario subject to external introgression. (A) Case of a 4-population tree subject to introgression from an external population *H*_5_. Consider *H*_2_ being the population subject to introgression from *H*_5_. (B) The subtree *T*_1:4_ includes the 4-population tree excluding the admixing population. (C) The subtree *T_out_* replaces the admixed population with the population source of introgression. (D) The subtree *T_un_*, where 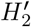 represents *H*_2_ when it has not yet undergone admixture, reflects the null hypothesis of correctness for the genetic relationship between four populations.

Consider the patterns of four alleles for the three subtrees mentioned above, whose estimated probabilities are respectively denoted as *p*_1:4_, *p_out_* and *p_un_*. Using the frequency estimators of eq (5) it is possible to estimate *p*_1:4_ and *p_out_*, but not *p_un_* since 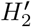 is not an observed population.

Assume that testing with the D-statistic on the tree *T*_1:4_ leads to a rejection of *H*_0_ because the allele frequencies of *H*_2_ are altered by the gene flow from *H*_5_. In fact, any combination of four alleles observed in T1 4 has probability

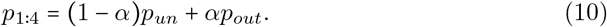

By solving for *p_un_* it follows that

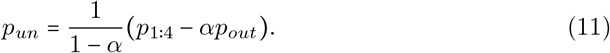

Note that if the admixture proportion *α* is known, then admixture correction is possible. If *α* is not known and we assume the tree is accepted for 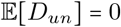, where *D_un_* is the D-statistic related to the tree *T_un_*, then *α* can be estimated. In this case, *p_un_* has to be determined for all values of *α*, and the correct one will be the value for which 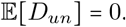. In this way an estimate of the admixture rate is obtained for the topology of Supplementary Figure 8A.

## Simulations

Different scenarios have been generated using `msms` [20] to reproduce the trees of Fig 2A, Fig 2B and Fig 2C, in which times are in units of generations. Each topology has been simulated 100 times for a constant population size of *N_e_* = 10^4^. Mutation and recombination of the simulations are consistent with human data [20]. Migrations and admixtures, respectively, for the scenarios of Fig 2A and Fig 2C, were simulated with specific options of `msms`. For each simulation we generated 200 regions of size 5MB for each individual and considered only variable sites, except for the case of Fig 2B, where the null hypothesis is affected by type-specific error on some of the individuals. We used a type-specific error of *e*_*A* → *G*_ = 0.005 in populations *H*_1_, *H*_3_. The choice of the region size is compatible with the one estimated in [12] to be sufficient to correct for the non independence of loci when bootstrapping with the jackknife estimator [21].

As a second step, the simulated genotypes from `msms` were given as input to `msToGlf`, a tool that is provided together with `ANGSD`. Using `msToGlf` it is possible to simulate NGS data from `msms` output files by generating the `pileup` files; that are used as input for `ANGSD`. As parameters for `msToGlf`, we set up the depth as mean of a poisson distribution and we hardcoded the error rates in the program when necessary for the scenario of Fig 2B.

**Fig 2.**
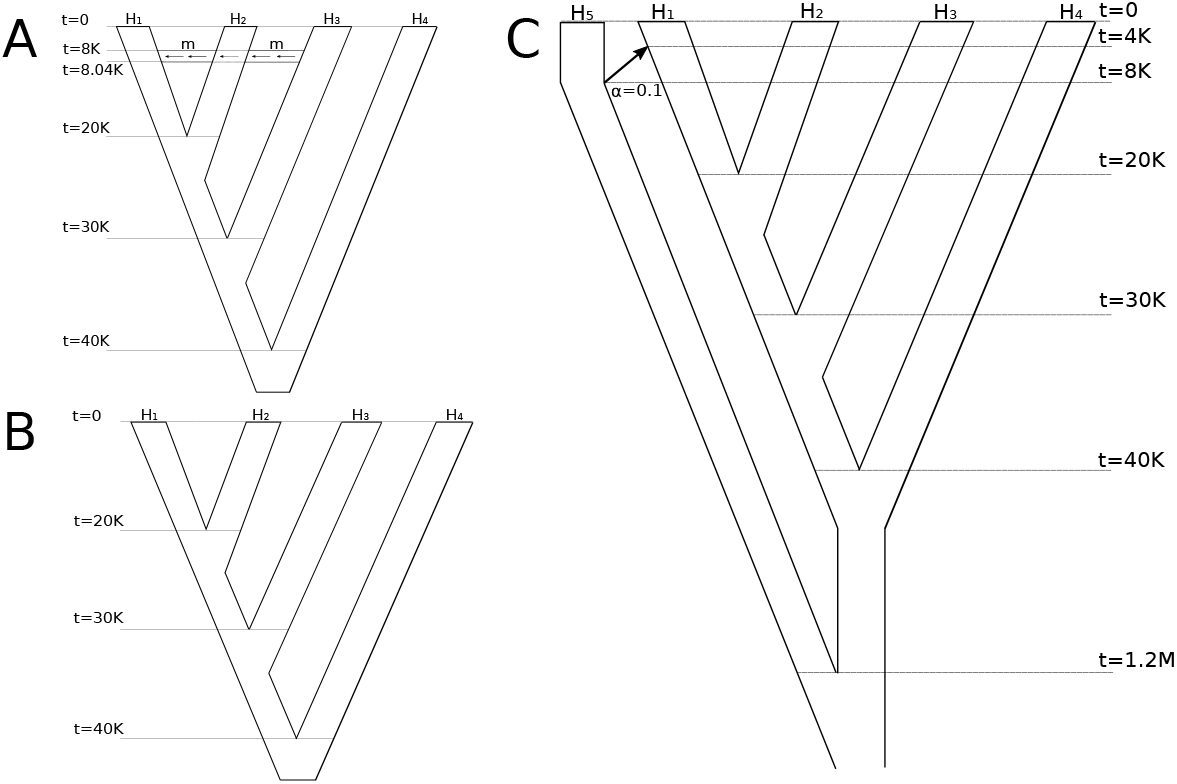
Simulated Scenarios. (A) Simulation of a tree in which migration occurs from population *H*_3_ to *H_i_*. The variable m is the (rescaled) migration rate varying between 0, 8, 16, 24, 32, 40 up to 280 with steps of size 20. Expressed in percentage, the migration rate varies between 0%, 0.02%, 0.04%, 0.06%, 0.08%, 0.1% up to 0.7%. Command: `msms -N 10000 -ms 40 200 -I 4 10 10 10 10 0 -t 100 -r 100 1000 -em 0.2 3 1 $m -em 0.201 3 1 0 -ej 0.5 1 2 -ej 0.75 2 3 -ej 13 4`. The same command line has been applied with the option `-I 4 40 40 40 40 0` to generate populations of 20 diploid individuals, used to study the power of the method using subsets of 1, 2, 5,10, 20 individuals of such populations. (B) Simulation of a tree in which no migration occurs, but type-specific errors on some individuals provide a rejection when testing for correctness of the null hypothesis. Command: `msms -N 10000 -ms 8 200 -I 4 2 2 2 2 0 -t 100 -r 100 1000 -ej 0.5 1 2 -ej 0.75 2 3 -ej 13 4`. (C) Simulation of a tree in which *H*_5_ admix with *H*_i_ with an instantaneous unidirectional admixture of rate *α* = 0.1. In this case we expect the null hypothesis to be rejected since *H*_5_ will alter the counts of ABBA and BABA patterns, but the alternative hypothesis does not involve gene flow with *H*_3_. Command: `msms -N 10000 -ms 50 200 -I 5 10 10 10 10 10 0 -t 100 -r 100 1000 -es 0.1 1 0.9 -ej 0.2 6 5 -ej 0.25 1 2 -ej 0.5 2 3 -ej 0.75 3 4 -ej 30 4 5`.

## Sequenced human populations

For the real data scenarios of Fig 3A, Fig 3B and Fig 3C we used Illumina sequenced individuals from several human populations. See Table 1 for an overview of the data. The depth of each individual has been calculated using the program `doDepth` of `ANGSD`. The Peruvian individuals used in our study were unadmixed with proportion ≥ 0.95. Estimation of the admixture proportions of these individuals was performed using `ADMIXTURE` [11]. In every individual, only the autosomal regions of all individuals were taken into consideration and bases were filtered out according to a minimum base quality score of 20 and a mapping quality score of 30. Type-specific error estimates for the Saqqaq, Mi’kmaq and French individuals were performed using the program `doAncError` of `ANGSD`, where the Chimpanzee was used as outgroup and the consensus sequence of human NA12778 as error-free individual (See Supplementary Figure 9 for the barplot of the estimates of the type-specific error).

**Fig 3.**
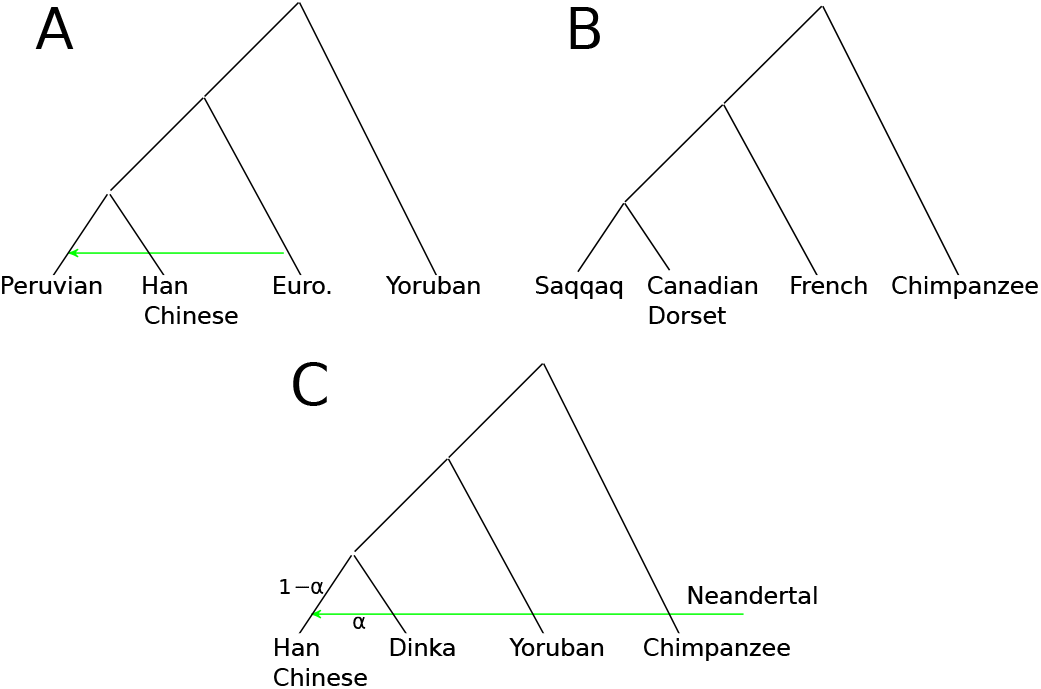
Real Data Scenarios. (A) Tree representing the southwestern European migration into the Americas during the European colonization. (B) Tree representing two independent migrations into northwestern Canada and Greenland. (C) Tree representing the presence of Neandertal genome into a modern non-african population, specifically the Han Chinese.

**Figure 9.**
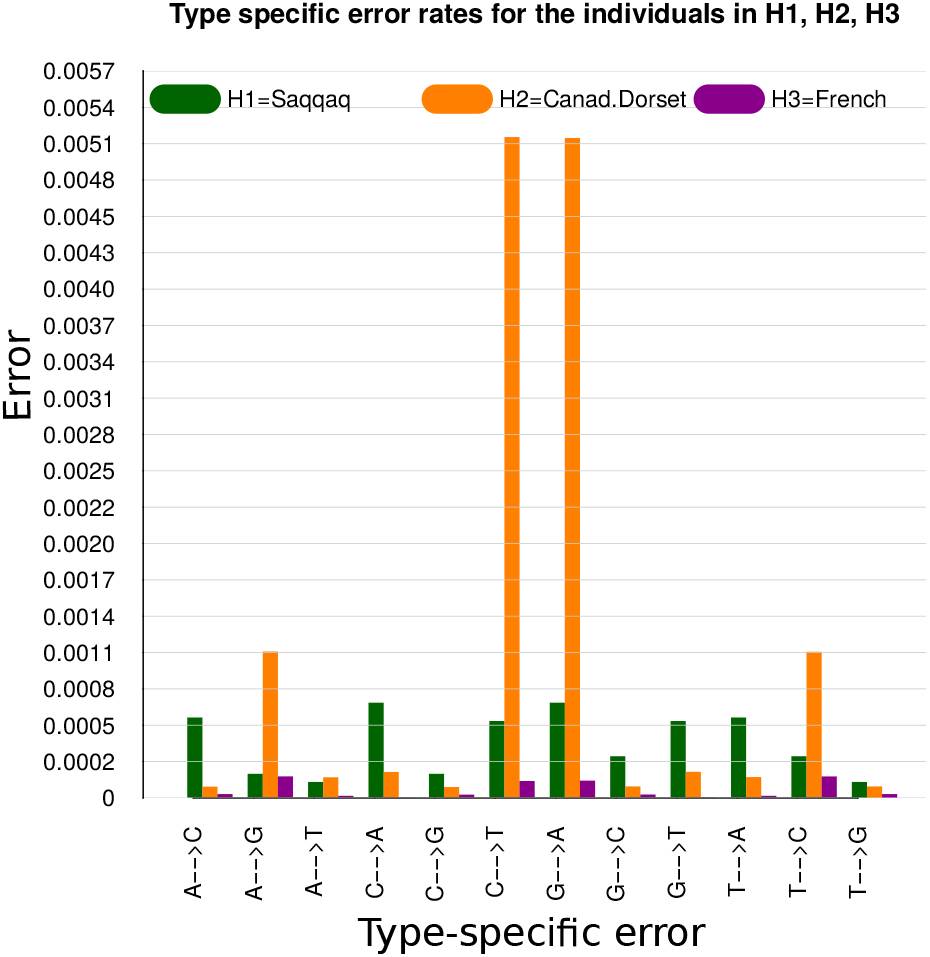
Estimates of Type-Specific Errors for Ancient Genomes. Estimated type-specific error rates for the Saqqaq, Mi’qmak and French genomes of the real data scenario illustrated in Fig 4B.

**Table 1.**
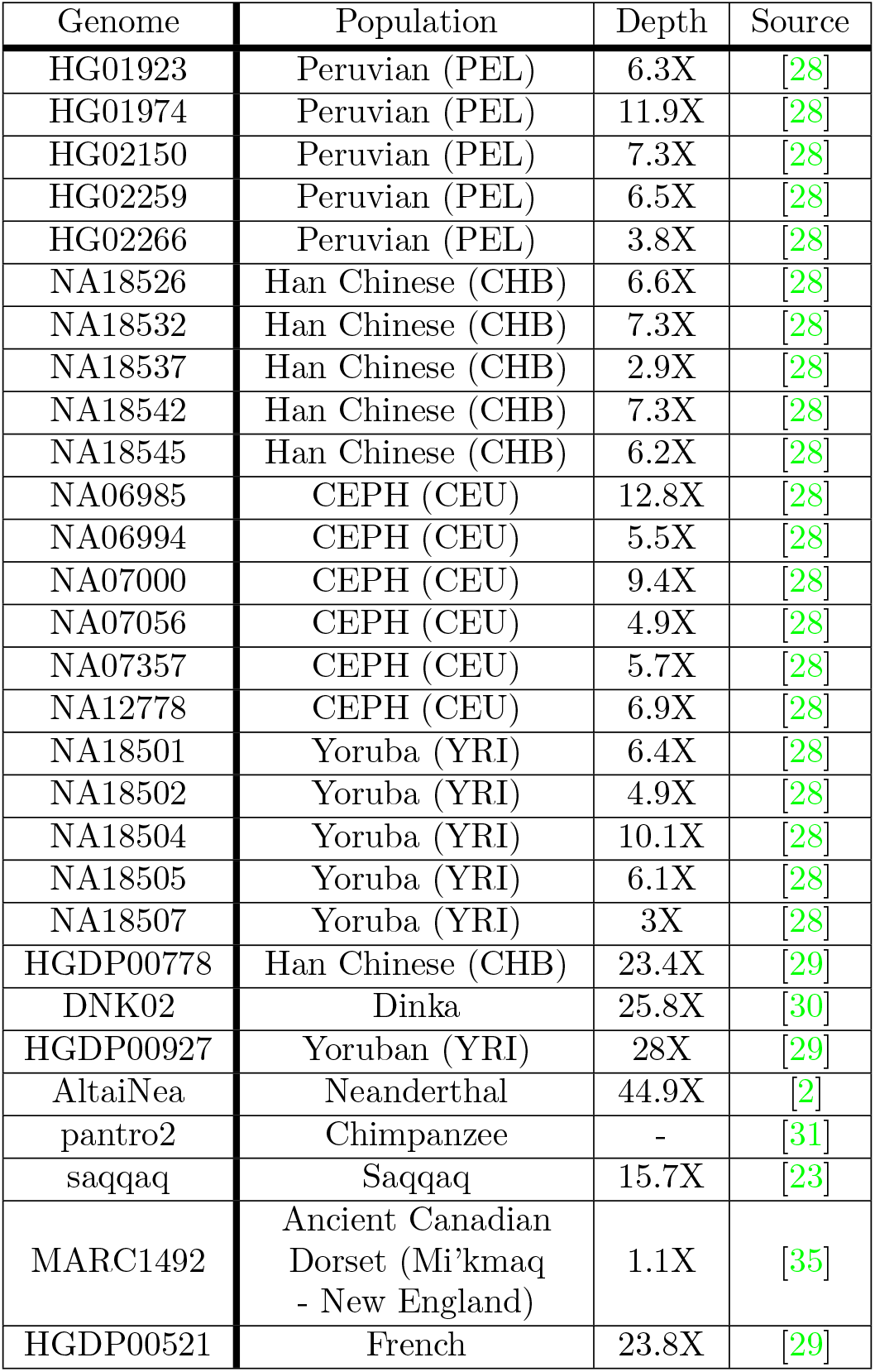
List of the Genomes Used in Real Data Scenarios. The table contains the genome identification number, the major population division, the depth calculated using ANGSD and the study source of the data.

## Results and Discussion

In the study of our results we compare different implementations of the D-statistic on simulated and real scenarios. We briefly define as *D_ext_* the extended D-statistic that we implemented, *D_ibase_* the D-statistic calculated by samplying 1 sequenced base per locus [2] and *D_geno_* the D-statistic calculated with equation (4) using the allele frequencies estimated from the true genotype (the true genotype is only available in the case of simulated data).

The D-statistic is computed on blocks of 5Mb, to ensure that every block is not subject to linkage disequilibrium from the other blocks, and that the number of loci in each block is large enough to make the D-statistic approach the approximation by a standard normal distribution (see Appendix 1). The use of blocks allows for estimation of a proper normalization constant for the D-statistic using the m-block jack-knife method [21]. The threshold for rejection of the null hypothesis is set to a p-value 0.001, corresponding approximately to the two-tailed acceptance region [-3,3].

The formula for calculating the D-statistic is given in eq (4) and finds amongst its current implementations, the ones in [15] and [16], with sampling of one base per locus from only one individual in each population. Such an implementation is computationally fast but has many drawbacks:

- when genomes are sequenced at low or medium depth (1X-10X), sampling one base might lead to a process with high uncertainty;
- base transition errors might affect the sampling of the base adding more uncertainty;
- only one individual per population is used;
- for a chosen individual chosen from a population, the reads are not used to evaluate the D-statistic, but only to sample one base.

We have proposed a solution to these problems with the extended version of the D-statistic *D_ext_* implemented in `ANGSD` and we will show in the following results how all the problems mentioned above are addressed.

## Comparison of Power Between the Different Methods

Using simulated and real data we compare the different types of D-statistics to study their sensitivity to gene flow, and illustrate how the improved D-statistic *D_ext_* is not affected by the issues faced by the current D-statistic *D_ibase_*, and even reach the performances of the D-statistic based on true genotype *D_geno_* at a rather low sequencing depth.

To evaluate the power of the different methods we first simulated NGS data based on coalescent simulations with mutation and recombination rates consistent with human populations [20]. We simulated without sequencing error four populations with a varying amount of migration from *H*_3_ to *H*_1_ (see Fig 2A) and applied the D-statistic based on five individuals from each population for two different sequencing depths. Fig 4A and Fig 4B show the power of the methods for depth 0.2X and 2X. Here power is the rejection rate of the null hypothesis when there is a migration from *H*_3_ to *H*_i_ in the tree (((*H*_1_,*H*_2_)*H*_2_)*H*_4_).

**Fig 4.**
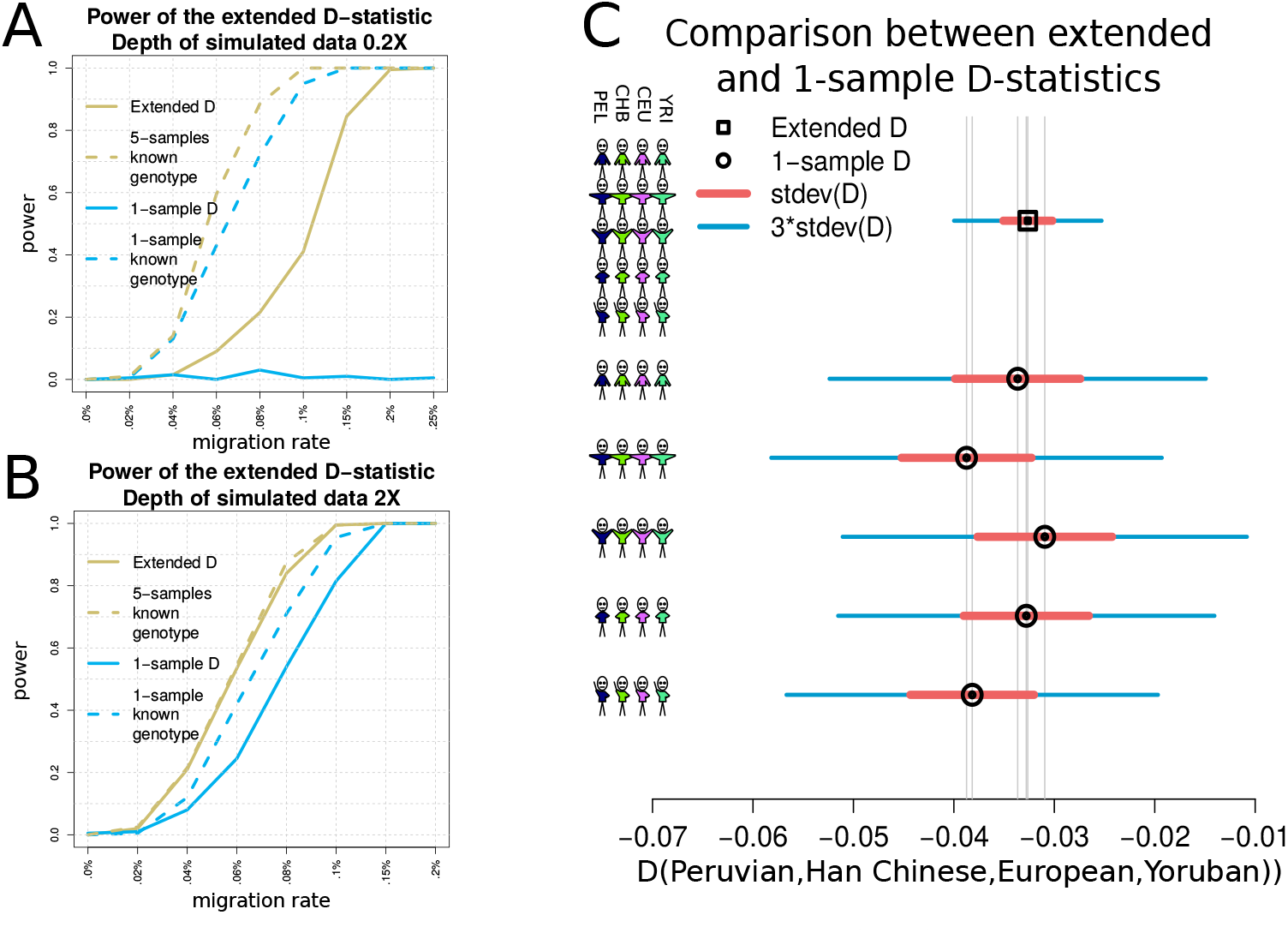
Detection of Admixture and Migration. (A,B) Rejection rate of the null hypothesis as a function of the migration rate in the tree (((*H*_1_,*H*_2_)*H*_3_)*H*_4_), where a migration from *H*_3_ to *H*_1_ occurs. The yellow and blue solid lines represent respectively the power of the method related to *D_ext_* and *D*_1*base*_. The yellow dashed line represents the rejection rate when the genotypes of the 5 individuals in each population are known and thus eq (4) can be applied. The blue dashed line illustrates the power of the method when only one genome per population has known genotypes. *D_ext_* performs almost as well as knowing the true genotypes already with depth 2X. (C) Value of *D_ext_* (black square) and values of *D*_1*base*_ (black circles) using respectively 5 genomes per population and one of them from each population. Each D statistic shows its associated standard deviation multiplied by 1 and 3. On the left side of the graph, the stickmen represent for each column the composition of the group by number of individuals.

The extended D-statistic proves to be effective in detecting gene flow even when the simulated depth is very low. For the scenario with sequencing depth 0.2X, *D_ibase_* is not able to detect almost any case of migration from *H*_3_, while *D_ext_* reacts with an acceptable rejection rate already for a migration rate as low as *m* = 0.15%. Of course such a very low depth does not allow the D-statistic to perform as well as *D_geno_*. In the case of sequencing depth 2X, *D_ibase_* does not always detect the alternative hypothesis and has also a considerable delay in terms of the migration rate necessary to do that, when compared to *D_ext_*. Furthermore *D_ext_* follows almost exactly the behaviour of the power related to *D_geno_*. This means that with a depth above 2X we can expect the D-statistic *D_ext_* to perform as well as knowing the exact genotypes of the data.

A deeper analysis to study the effect of using multiple individuals per group is illustrated in supplementary Figure 7. Here we simulated again the scenario with depth 0.2X, and compared the use of 1, 2, 5, 10 and 20 individuals per population. The graph shows that using multiple individuals increases the power of the method and at the same time decreases the standard deviation of *D_ext_*.

**Figure 7.**
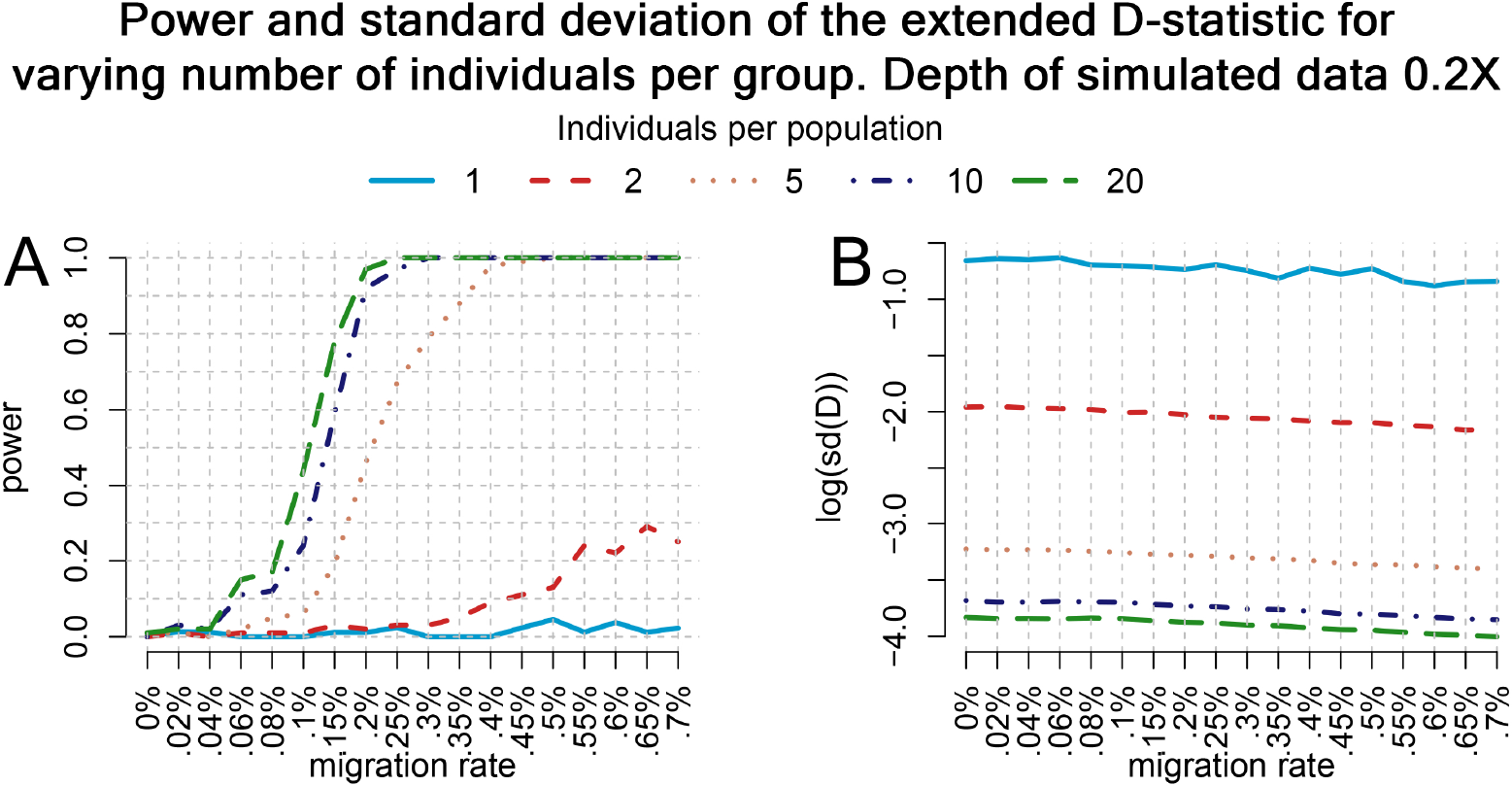
Effect of the number of individuals per population in detecting admixture. Results from the simulation of the scenario of Figure 2A, subject to a migration from *H*_3_ to *H*_1_, using either 1, 2, 5, 10 or 20 individuals per population sequenced at depth 0.2X. (A) Power of the extended D-statistic for increasing values of the number of individuals per group. (B) The value of the standard deviation of *D_ext_* for different number of individuals per population.

The power of *D_ext_* and *D_ibase_* are compared in a real data scenario using Illumina sequenced modern human populations from the 1000 Genomes Project with a varying sequenced depth in the range 3-13X. We specifically used PEL=Peruvian, CEU=European, CHB=Han Chinese and YRI=African Yoruban individuals to form the tree (((PEL,CHB)CEU)YRI) shown in Fig 3A. This scenario represents the southwestern European gene flow into the ancestors of the Native Americans [13]. Each of the four populations consists of 5 sequenced individuals when evaluating *D_ext_*, and a distinct one of those individuals when evaluating *D_ibase_* five times (see Fig 4C). The extended D-statistic *D_ext_* has much lower standard errors, that corresponds to a smaller p-value than in the case of *D_ibase_*, and therefore a more significant rejection. See Supplementary Table 2 for a better comparison of the values of the different D-statistics.

**Table 2.** European Introgression into Native American Individuals. The table contains the values of the different types of D-statistics used to create the plot of Fig 5C, reporting the D-statistic for the tree (((PEL,CHB)CEU)YRI). The first column denote if we are illustrating either the extended D-statistic, *D_ext_*, or the D-statistic that uses a sampled base, *D*_1*base*_. The column denoted by D is the D-statistic over all blocks of loci, used to estimate the standard deviation (third column) by bootstrapping. The Z-score represents the D-statistic normalized by its standard deviation. The last column represents the ratio between the estimated standard deviations of *D*_1*base*_ and *D_ext_*.

| D-statistic | D | stdev(D) | Z-score | $\frac{\sigma_{1base}}{\sigma_{ext}}$ |
| --- | --- | --- | --- | --- |
| $D_{ext}$ | -0.032638 | 0.002449 | -13.114101 | - |
| $D_{1base}$ | -0.038171 | 0.006164 | -6.223641 | 2.51 |
| $D_{1base}$ | -0.032786 | 0.006244 | -5.253267 | 2.54 |
| $D_{1base}$ | -0.030950 | 0.006708 | -4.602315 | 2.74 |
| $D_{1base}$ | -0.038730 | 0.006480 | -5.999972 | 2.64 |
| $D_{1base}$ | -0.033640 | 0.006244 | -5.353646 | 2.55 |

It is worth to underline that the presence of structured populations might lead to false positives because the structure is not considered in the model. If there is structure within *H*_1_,*H*_2_, the properties of the D-statistic are preserved. However, if the population was structured prior to the split of H1 and H2, then it will affact the D-statistics.

## Error Impact and Correction

Sequencing or genotyping errors are known to have a large impact on the D-statistic [19]. Using simulation we show that if the type-specific error rates are known then we can correct the D-statistic accordingly. We simulate the tree under the null hypothesis. However, we add base *A* → *G* error rate of 0.005 in populations *H*_1_ and *H*_3_ in order to alter the observed number of ABBA and BABA combination of alleles, and consequently lead to a possible rejection of the null hypothesis.

In the plot of Fig 5A are represented the estimated distributions of the Z-scores related to *D_ext_* before and after error estimation and error correction, for 100 simulations of a tree (((*H*_1_,*H*_2_)*H*_3_)*H*_4_) without any gene flow, where we have also introduced type-specific error for transitions from allele A to another allele for the individuals in *H*_1_,*H*_2_,*H*_3_ at different rates. The test statistic has high values due to the error while all simulations fall in the acceptance interval if we perform error correction.

**Fig 5.**
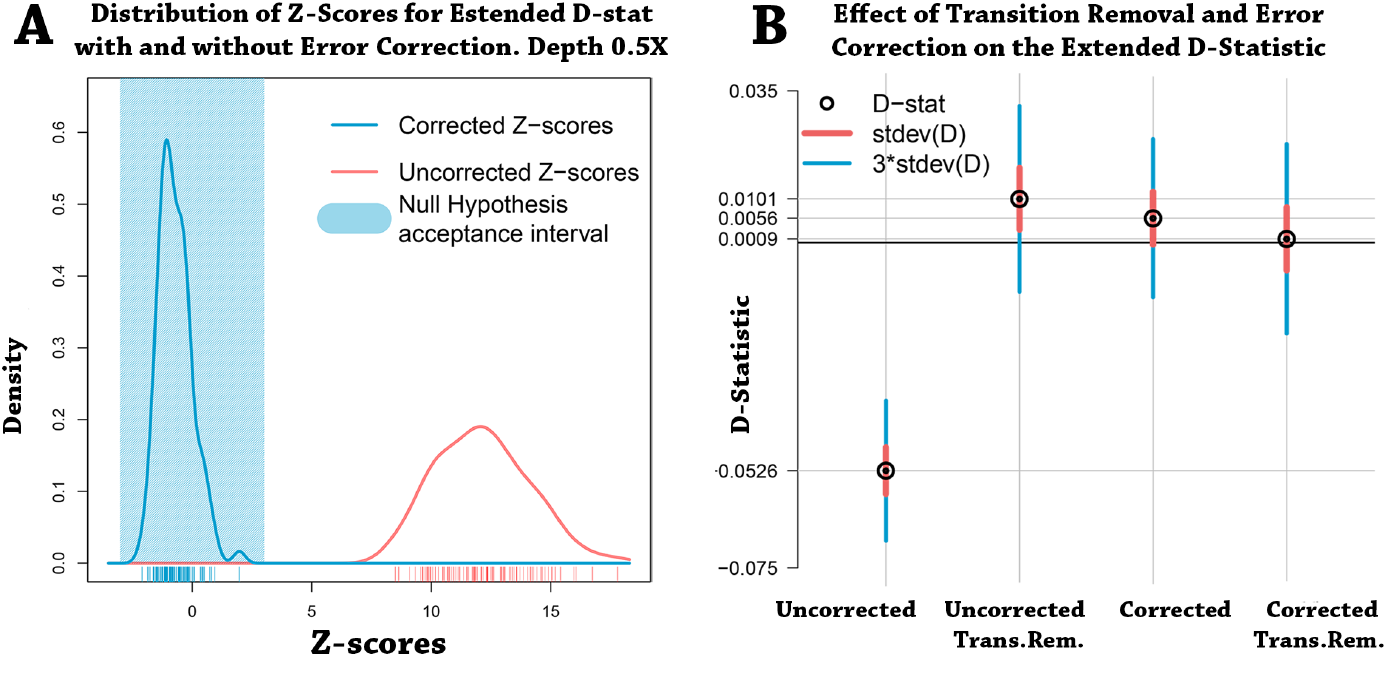
Effect of Error Estimation and Correction. (A) Estimated distributions of the Z-scores related to *D_ext_* for the null hypothesis (((*H*_1_, *H*_2_)*H*_3_)*H*_4_) in which *H*_1_,*H*_3_ and *H*_2_ has probability 0.005 and 0.01 of transition from base A, respectively. The blue polygon represents the interval where a Z-score would accept the null hypothesis. The red line represents the distribution of Z-scores before type-specific errors are corrected. In blue we have the Z-scores after correction. (B) Values of *D_ext_* in four different cases for the tree (((Saqqaq, Dorset)French)Chimpanzee). The black circles are the values of the uncorrected D-statistic, removal of ancient transitions, error correction, error correction and ancient transitions removal. The red and blue lines represent the standard deviations and the value they need to reach the threshold of |*Z*| = 3, respectively.

The uncorrected D-statistic performs poorly because of the errors in the data that cause rejection of the null hypothesis in all simulations. It is remarkable to observe that *D_ext_* has good performances already at depth 0.5X. This means that even small error rates in the data make the D-statistic very sensible to the rejection of *H*_0_. Therefore we require to apply error correction to our data. The result is that the Z-scores fall into the acceptance threshold and the null hypothesis is fulfilled. The distribution of corrected Z-scores is not perfectly centered in 0 because of imperfect error correction.

The most obvious need for error correction in real applications is the use of ancient genomes, which have a large amount of errors, especially transitions. To illustrate the effect of errors in real data and our ability to correct for them we use two ancient genomes which contain a high sequencing error rate due to *post mortem* deamination. The tree (((Saqqaq,Dorset)French)Chimpanzee) of Fig 3B illustrates the migrations to western Canada (Canadian Dorset Mi’kmaq genome) and southwestern Greenland (Saqqaq genome). Due to the effect of deamination prior to sequencing [23,35], the two ancient genomes have high type-specific error rates as shown in Supplementary Table 3 and Supplementary Figure 9. The error rates alter the counts of ABBA and BABA patterns, which bias the uncorrected D-statistic.

**Table 3.** Estimated Error Rates. Estimated type-specific error rates for the ancient individuals Saqqaq and Canadian Dorset Mi’kmaq used in the tree of Figure 3B.

| Individual | $A \rightarrow C$ | $A \rightarrow G$ | $A \rightarrow T$ | $C \rightarrow A$ | $C \rightarrow G$ | $C \rightarrow T$ |
| --- | --- | --- | --- | --- | --- | --- |
| Saqqaq | 1.90e-04 | 6.08e-04 | 3.27e-04 | 7.52e-04 | 1.22e-04 | 6.32e-04 |
| Dorset | 8.86e-05 | 1.15e-03 | 1.62e-04 | 2.04e-04 | 8.52e-05 | 5.22e-03 |
| | $G \rightarrow A$ | $G \rightarrow C$ | $G \rightarrow T$ | $T \rightarrow A$ | $T \rightarrow C$ | $T \rightarrow G$ |
| Saqqaq | 6.35e-04 | 1.26e-04 | 7.52e-04 | 3.28e-04 | 6.08e-04 | 1.91e-04 |
| Dorset | 5.21e-03 | 9.01e-05 | 2.06e-04 | 1.64e-04 | 1.15e-03 | 9.04e-05 |

We expect the tree to be true under the null since Saqqaq and Dorset have a recent common ancestor [22]. In Fig 5B we compare the extended D-statistic *D_ext_* in four cases: firstly using observed data, secondly removing all transitions which are related to most of the errors, thirdly applying error correction and lastly combining error correction and transitions removal. Note that the removal of transitions related to the pairs of alleles A,C and G,T is the current standard technique to avoid high error rates when calculating the D-statistic from damaged low-coverage data. The uncorrected D-statistic rejects the null hypothesis whereas correction or transition removal gives a non-significant test. Error correction performs better than transition removal, providing a value of the D-statistic that is closer to 0 and has smaller standard deviation. Supplementary Table 4 shows the values related to the four D-statistics in this scenario. Supplementary Figure 10 illustrates the effect of increasing and decreasing the removal of error for the base transition *C* → *G* and *C* → *T* for one of the Saqqaq, Dorset and French genomes. This correspond to add a value to the estimated error rate matrix of one of the individuals. Observe that the French individual is less affected by the addition or removal of error than the first two individuals. Moreover all 3 individuals are more sensible to the error rate in the case of transversion *C* → *T*.

**Figure 10.**
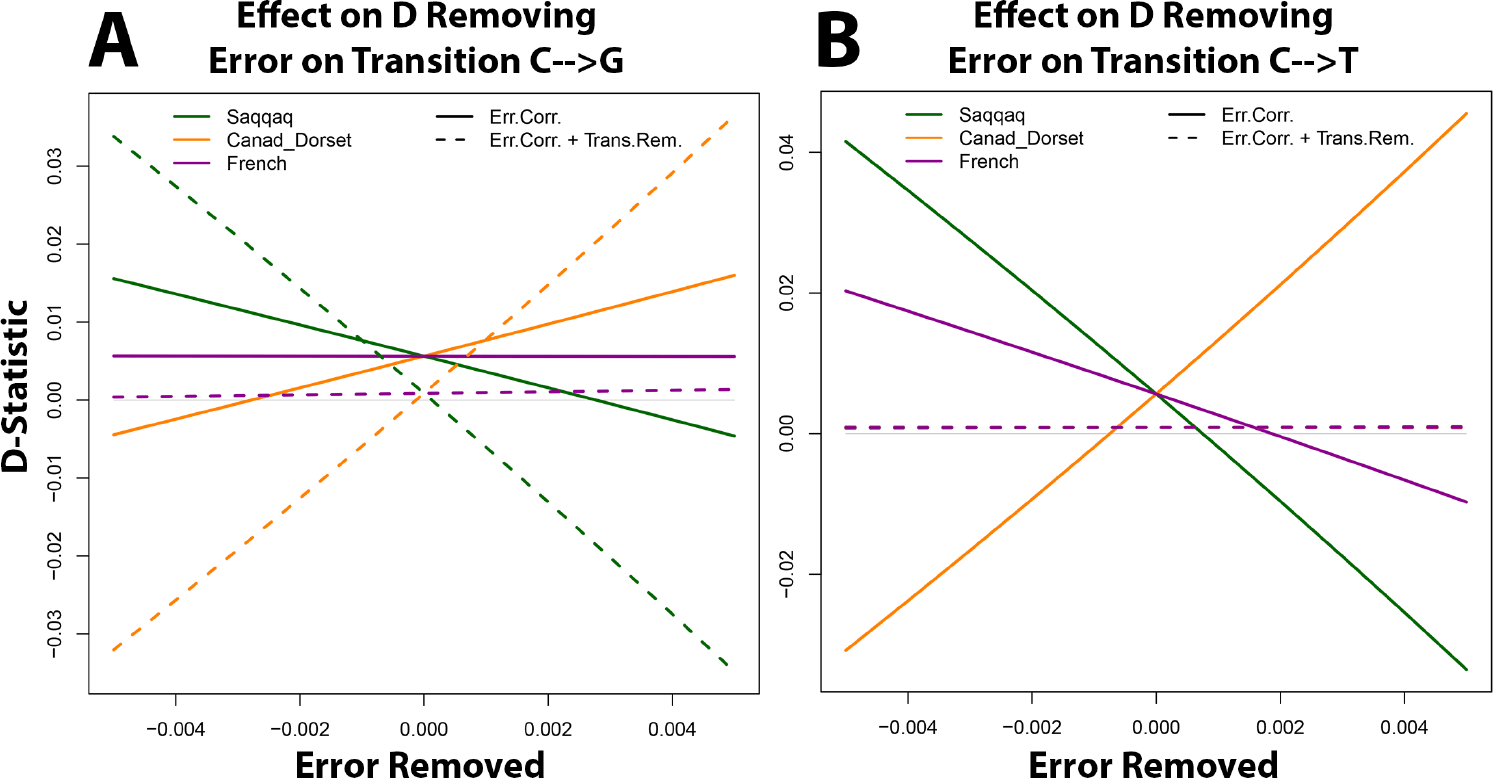
Behaviour of the D-Statistic in Function of the Type-Specific Error. Effect of increasing and decreasing the removal of error for the base transitions *C* → *G* and *C* → *T* for one of the Greenlandic Saqqaq, Canadian Dorset and French genomes. This corresponds to the addition of a value in the entry ***e**(G,C)* or ***e**(T, C)* of the estimated error matrix of one of the individuals, as if the estimated error rate was higher or lower. In solid lines are represented the values of *D_ext_* for which the correction is performed. The dashed lines represent the analogous values where ancient transitions are not considered.

**Table 4.** Extended D-Statistic in Real Data Scenario with Ancient Genomes. Table comparing the extended D-statistic with the application of error correction and/or transition removal for the tree of Figure 3B, where the ancient individuals Saqqaq and Canadian Dorset Mi’kmaq are affected by high type-specific error rates.

| Correction | $D_{ext}$ | $sd(D_{ext})$ | $Z - score$ | $p - value$ |
| --- | --- | --- | --- | --- |
| None | -5.26e-2 | 5.4e-3 | -9.81 | 0 |
| Trans.Rem. | 1.01e-2 | 7.1e-3 | 1.41 | 1.57e-1 |
| Error.Corr. | 5.64e-3 | 6.1e-3 | 0.93 | 3.51e-1 |
| Err.Corr & Tr.Rem | 8.77e-4 | 7.3e-3 | 0.12 | 9.04e-1 |

## Correction for External Introgression

We use simulations of a scenario with external introgression to verify the performance of correction for gene-flow in restoring a four-population tree configuration that lead to the acceptance of the null hypothesis *H*_0_. In the simulation case we know the value of *α*, that is the amount of introgression, therefore correction is possible. Thereafter we use a known genetic relationship involving the Neandertal introgression into out-of-Africa modern individuals in Europe and Asia [2,4] to correct for the effect of admixture. In addition we show that, if we assume the absence of gene flow in the tree topology, then we can estimate the amount of introgression, and compare it with the estimation involving the original D-statistic tools.

For some species there are introgression events from an external source which can affect the D-statistic when performing test for admixture among the species. We performed 100 simulations of the null hypothesis (((*H*_1_,*H*_2_)*H*_3_)*H*_4_) of Fig 2C, for which an external population *H*_5_ is admixed with *H*_2_ with rate *α* = 0.1. The plot of Fig 6A shows the estimated distribution of the Z-scores related to the observed and admixture-corrected *D_ext_*. The observed D-statistic is positive and has Z-scores that reject the null hypothesis. Applying eq (11) we are able to remove the effect of gene flow from *H*_2_. The result of removal of the gene flow’s effect is that the estimated probabilities of ABBA and BABA combinations of alleles are altered and the resulting calculated values of the D-statistic lead to acceptance of the null hypothesis *H*_0_.

**Fig 6.**
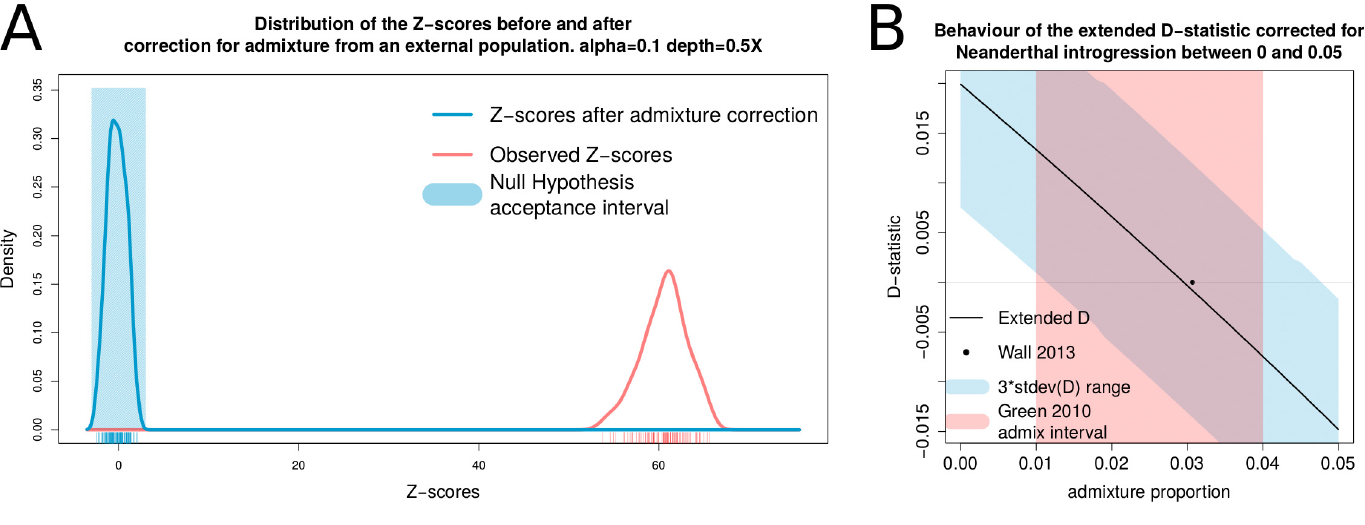
Effect of Correction from External Introgression. (A) Estimated distribution of the Z-scores related to *D_ext_* from the 100 simulations of the null hypothesis (((*H*_1_, *H*_2_)*H*_3_)*H*_4_) with introgression of rate *α* = 0.1 from an external population *H*_5_ into *H*_2_. The Z-scores of the observed tree are far off the acceptance interval because of the admixture from *H*_5_. Once the portion of genome from the external population is removed from *H*_2_, the tree fulfills the null hypothesis and the Z-scores all fall in the acceptance interval defined by |*Z*| ≤ 3. (B) Behaviour of the *D_ext_* of the tree (((Han Chinese,Dinka)Yoruban)Chimpanzee) as a function of the admixture rate a used to correct for the introgression of the Neandertal population into the Han Chinese population. The red polygon is the interval in which [2] estimates *a* to fall in. The black dot coincides with the value of *α* = 0.0307 calculated by [4] using the tree (((Han Chinese,Yoruban)Neandertal)Chimpanzee), with standard deviation 0.0049. The blue polygon is 3 times the standard deviation of *D_ext_*. When *D_ext_* is 0, we estimate *α* = 0.03 with standard deviation 0.0042.

For human populations it is problematic to use the D-statistics when applied to both African and non-African populations because of ancient gene-flow from other hominids into non-Africans. Therefore, *H*_0_ might not fulfilled for any tree (((*H*_1_,*H*_2_)*H*_3_)*H*_4_) where an ingroup consists of both an African and a non-African population. This leads to rejection of the tree and to the natural conclusion that there is gene flow between *H*_3_,*H*_2_ (resp. *H*_3_,*H*_1_). However, if there is known external admixture from a population *H*_5_, it is possible to correct for admixture from this external contribution.

We illustrate the problem and our ability to correct for it using the tree shown in Fig 3C, which shows introgression of the Neanderthal genome into the ancestors of the Han Chinese population. The correction is performed for the admixture proportion *α* in the range [0, 0. 05] in steps of 0. 01. The value of *α* for which the *D_ext_* is closest to 0 might be considered as an estimate of the admixture rate. We choose these populations because we can compare our result with the estimate from previous studies of the same populations [2,4]. Green et al. [2] estimated *α* to be in the range [0.01,0.04], while Wall et al. [4] estimated it as being *α* = 0.0307 with standard deviation 0.0049. The result is shown in Fig 6B for the tree (((Han Chinese,Dinka)Yoruban)Chimpanzee) for different admixture rates *α* used to correct for the introgression of the Neandertal population into the Han Chinese population. The red polygon is the interval in which *α* is estimated to be [2]. The black dot coincides with the value of *α* = 0.0307 calculated in [4]. The blue polygon is 3 times the standard deviation of *D_ext_*. For almost the whole range of reported admixture proportions, the tree is not rejected after adjustment for admixture, indicating that the uncorrected D-statistic concluded the presence of gene flow. When *D_ext_* is 0, we estimate *α* = 0. 03 with standard deviation 0.0042, which is similar to previous estimates.

In both the cases of simulated and real data we have thus been able to distinguish the case in which the alternative hypothesis is due to an external introgression and not to admixture from *H*_3_. In our simulations, the admixture correction seems not to suffer from the effect of drift, which is not modelled in the correction. In fact the branch leading to *H*_5_ splits 8000 generations in the past and admixes 4000 generations in the past on the branch leading to *H*_1_. Thus there is a drift affecting gene frequencies of both the admixing and admixed populations.

In the case of real data the exact amount of admixture *α* is not previously known. Therefore we calculated the D-statistic for the tree (((Han Chinese,Dinka)Yoruban)Chimpanzee) using admixture-corrected values of the probabilities of allele patterns, considering values of the admixture rate falling in the interval estimated in [2]. Without admixture correction, the obvious conclusion would have been that for the tree (((Han Chinese,Dinka)Yoruban)Chimpanzee) there is gene flow between the Yoruban and Dinka populations.

## Conclusions

In summary we have implemented a different D-statistic that address the drawbacks of the current implementations of the D-statistic, but still preserve the approximation as a standard normal distribution (see Appendix 1) that allows for a statistical test. The extended D-statistic D_ext_ allows for multiple individuals per population and instead of sampling one base according to the estimated allele frequencies, uses all the available sequenced bases.

Using both simulations and real data we have shown that

1. the extended D-statistic *D_ext_* has more power than the alternative methods, with an increased sensibility to admixture events;
2. the performance of the extended D-statistic is the same as when true genotype is known for a depth of at least 2X,
3. we can accomodate type-specific errors to prevent that en eventually wrong acceptance or rejection of the null hypothesis is caused by error-affected allele frequencies. The error estimation and correction reveal to be especially suited in the case of ancient genomes, where error rates might be high due to chemical treatments prior to sequencing and degradation over time;
4. we can calculate the D-statistic after correcting for admixture from an external known population, such as in the case of Neandertal gene flow into the Han Chinese population.

The extended D-statistic *D_ext_* is especially effective compared to the standard D-statistic *D_lbase_* when applied to data with low/variable depth, multiple individuals and ancient DNA.

## Appendices

The setup of the theoretical treatment consists of four sampled genomes representing four populations *H*_1_, *H*_2_, *H*_3_, *H*_4_, for which we assume the relationship illustrated in Fig 1. Each genome is considered to have *M* di-allelic loci. We will consider the situation in which *M* grows to infinity. Each locus *i* consists of a certain number 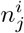 of alleles A and B, where *j* = 1, 2,3,4, is the index of the *j*th genome. Moreover we assume independence between the loci.

Assume that at a locus *i* the allele frequencies in the four groups of individuals 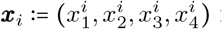 follow a locus-dependent distribution *F_i_*(***x***), *i* = 1,…, *M* and let 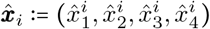 be an unbiased estimator of ***x***_*i*_ at locus *i*, such as the relative frequencies of the allele A in each population. The populations’ frequencies are considered to be a martingale process.

The null hypothesis that the tree of Fig 1 is correct can be rewritten as follow:

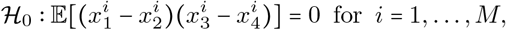

where the expectation is done on the difference between the probabilities of ABBA and BABA events deduced in eq (1) and eq 2. Using the empirical frequencies as proxies for the expected values, we build the following normalized test statistic, also known as D-statistic:

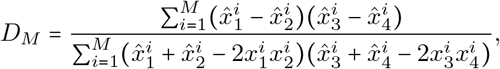

where the values

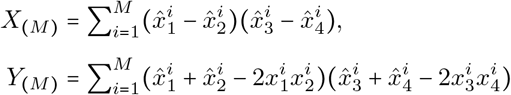

are the numerator and denominator of the D-statistic, respectively.

### Appendix 1 Convergence of the D-Statistic

In this paragraph we prove that the D-statistic defined as

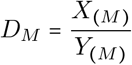

converges in distribution to a standard normal variable up to a constant.

Rewrite the numerator and denominator as

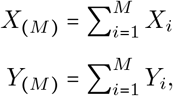

where the values *X_i_* and *Y_i_* are defined for each *i* = 1,…, *M* by

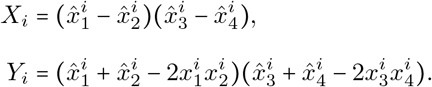

Consider the series of independent variables *X_i_* in the numerator of *D_M_*, having means *μ_i_*. Every term *X_i_* of the numerator is an unbiased estimate of 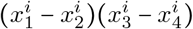, assuming the observed allele counts are binomially distributed [12]. We show in the following proposition that every term of the numerator of the D-statistic has expectation *μ_i_* = 0 for *i* = 1,…, *M* by calculating the expectation of 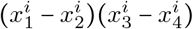.

**Theorem 1**. *Given the tree topology of Fig 1, it holds that* 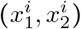 *for i* = 1,…,*M*.

*Proof.* Let 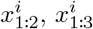 and 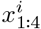 be the frequencies of the ancestral populations of 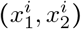, 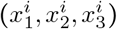 and the root of the tree, respectively, as illustrated in Fig 1. Let ***χ*** be the set of those three frequencies. Using the martingale properties of the frequencies it follows that

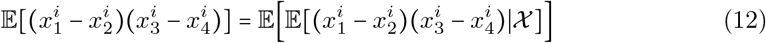

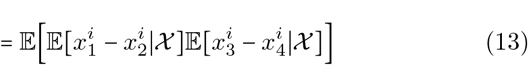

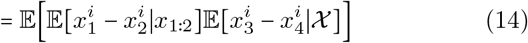

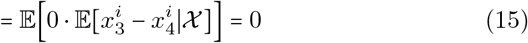

Therefore *X_i_* has mean 0 for all *i* = 1,…, *M*.

To prove convergence of the D-statistic for large *M* we assume the following:

1. Let 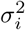 be the variance of every term *X_i_*. Denote with *v_M_* the sum 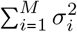, then

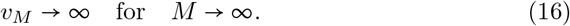
2. Let *Y_i_, i* = 1,…, *M*, be the series of independent variables in the denominator of *D_M_*, having means *γ_i_*. Then

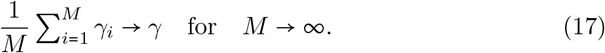
3. Denote with 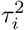 the variance of *Y_i_*. Then

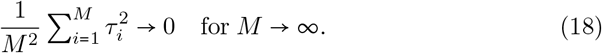

If the numerator and denominator are sums of iid variables, conditions (16), (17) and (18) are fullfilled. In fact, if every term *X_i_* has variance *σ*^2^, the sum of variances is *v_M_* = *Mσ*^2^ and eq (16) holds. If every term *Y_i_* has mean and variance *γ* and *τ*^2^, respectively, eq (17) is still valid because the arithmetic mean is done on identical values. Moreover, eq (18) holds because

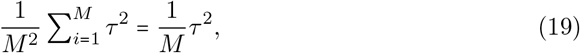

that converges to zero for *M* → ∞.

The convergence of the D-statistic *D_M_* is proved in steps, analyzing separately the numerator and the denominator. We begin by stating all the necessary theorems. Firstly, we consider an extension of the central limit theorem (CLT) [24], that will be applied to the numerator *X*_(*M*)_. Subsequently we state the law of large number (LLN) [25] for not iid variables that is used for the denominator *Y*_(*M*)_ of the D-statistic. Thereafter we enunciate one of the consequences of Slutsky’s theorem [26,27]. The last step is a theorem for the convergence of the D-statistic, proved by invoking all the previous statements, applied to the specific case of the D-statistic.

**Theorem 2** (CLT for independent and not identically distributed variables). *Let* 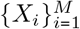 *be a sequence of independent (but not necessarily identically distributed) variables with zero mean and variances* 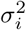. *Define v_M_ as* 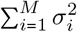. *Consider the following quantity*

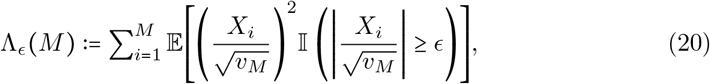

*where* II(·) *defines the indicator function. If for any ∊* > 0 *it holds that* lim_*M*→∞_ Λ_∊_(*M*) = 0, *then the normalized sum* 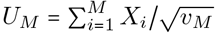 *converges in distribution to a standard normal **N***(0,1).

**Theorem 3** (LLN for independent and not identically distributed variables). *Let* 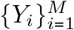 *be a sequence of uncorrelated random variables. Define* 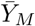 *as the empirical average* 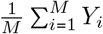. *Denote with γ_i_ and* 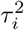 *the expectation and variance of each variable. If conditions* (17) *and* (18) *are fulfilled, then for each ∊* > 0

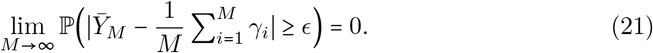

*Equivalently the empirical average* 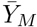 *converges in probability to* 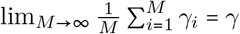.

**Theorem 4** (Slutsky’s Theorem). *Let X_(M)_ and Y_(M)_ be two sums of not iid random variables. If the former converges in distribution to X and the latter converges in probability to a constant γ for M* → ∞, *then the ratio X_(M)_/Y_(M)_ converges in distribution to X/γ*.

The last step is a theorem for the convergence of the D-statistic, proved by invoking all the previous statements, applied to the specific case of the D-statistic.

**Theorem 5** (Convergence in distribution of the D-statistic). *Consider the D-statistic defined by*

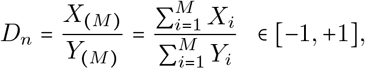

*where numerator and denominator are sum of independent (but not necessarily identically distributed) variables. Under the assumptions of* (16), (17) *and* (18), *the D-statistic converges in distribution to a standard normal if rescaled by the constant*:

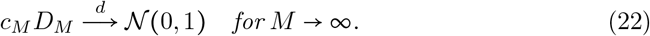

*The arrow denotes the convergence in distribution and CM is defined as*

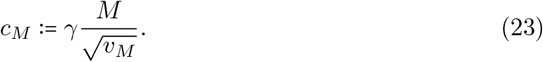

*Here v_M_ is the sum of the variances of the first M terms of the numerator, and γ is the convergence value of thee aritmetic mean of the denominator’s expectations for M* → ∞.

*Proof.* First consider Theorem 2 applied to the rescaled numerator 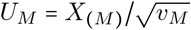. It is necessary to prove that for any *∊* > 0 it holds that lim_*M* →∞_ Λ_*∊*_(*M*) = 0 to ensure the convergence in distribution. First observe that |*X_i_*| ≤ 1 for any index *i*. Consequently we have the inequality

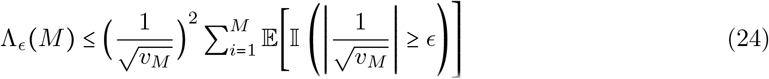

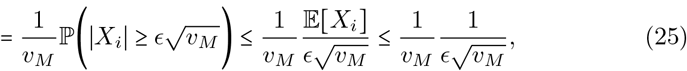

where Markov’s inequality is applied to the last line of the equation. Thus *U_M_* converges in distribution to a standard normal *N*(0, 1)

Since conditions (17) and (18) are fulfilled by assumption, it is possible to invoke Theorem 3 to state that the empirical average of the denominator *Y_(M)_/M* converges in probability to a constant *γ*, which is positive since every term of the denominator is positive.

Finally, we apply Theorem 4 using the proper constants that follows from Theorems 2 and 3 applied to the numerator and denominator, respectively. We proved that the sum 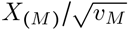 converges in distribution to a standard normal *N*(0,1) and *Y_(M)_/M* converges in probability to the constant *γ*, that is the limit of the arithmetic mean of eq 17. Thus the ratio

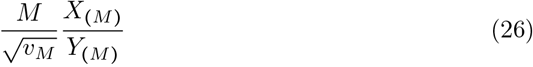

converges in distribution to a gaussian 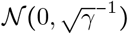. The convergence in distribution of *D_M_* to a standard normal variable is accomplished by rescaling by the following multiplicative constant

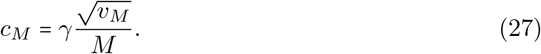

The results of this proof apply also in the following cases of the D-statistic:

1. the original D-statistic *D_M_* calculated by sampling a single base at each site from the available reads [2] to estimate the sampling probabilities. In this case every term on the numerator has possible values -1, 0, +1. Each population frequency xj is parameter of a binomial distribution 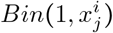, and is estimated by the frequency of the observed base A at locus *i* in population *j*,
2. the D-statistic is evaluated using the estimated population frequencies 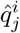 defined in eq 5 for multiple individuals in a population (see Appendix 2). In fact, the estimator for multiple individuals is still an unbiased estimate for the population frequency [18], therefore every term of the numerator is still an unbiased estimate for the difference between the probabilities of ABBA and BABA events.
3. the D-statistic is evaluated only over loci with allele frequency *x*_4_ = 1 for population *H*_4_. This special case of D-statistic has been used, for example, to assess the presence of gene flow from the Neandertal population into modern out-of-Africa individuals, setting a Chimpanzee as outgroup, and considering only loci where the outgroup showed uniquely allele A [2]. in fact, Theorem 1 still holds because in eq (12) the term 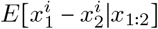 is zero, independently of which values 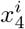 assumes.

### Appendix 2 Multiple Genomes

We assume a di-allelic model with alleles A and B and the four populations *H*_1_, *H*_2_, *H*_3_, *H*_4_ that consist each of a number of distinct individuals *N_j_, j* = 1,2,3,4, where *j* indexes the populations. Given the allele frequency 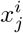, *j* = 1, 2, 3, 4, at locus *i*, we model the observed data as independent binomial trials with parameters 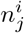 and 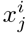 for *j* = 1, 2,3,4, where 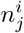 is the number of trials. One possible unbiased estimator of the population frequency is

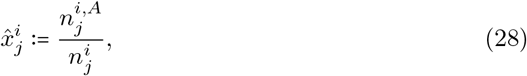

where 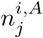 is the total number of As and 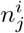 the total number of bases observed for the selected population and locus.

For locus *i* denote the allele frequency of individual *ℓ* in population *j* as 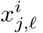. We use as its unbiased estimator

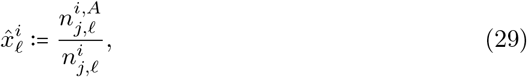

namely the ratio between the number of observed As and the total number of observed alleles at locus *i* in genome *ℓ*. The idea is to condense all the quantities 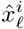 into a single value 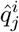 that minimizes the variance of the sum of the estimated individuals’ frequencies w.r.t. a set of normalized weights

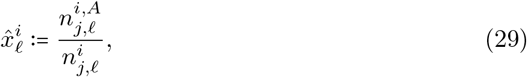

The estimated population frequency 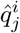 is an unbiased estimator of the frequency of population *j* at the ith locus [18]. The aim of the weight estimate is to determine the set of weights that minimizes the variance of 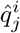. To do this, we first determine the variance of each individual’s frequency.

Consider a genome *ℓ* in population *j*. We approximate the frequency estimator of genome *ℓ* in population *j*, namely 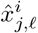, defining

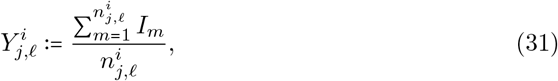

where 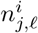 is the total number of reads for individual *ℓ* and 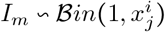 for 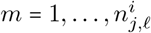. Note that the Binomial variables are parametrized by 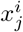 and not by 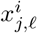. The variance of 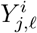 is

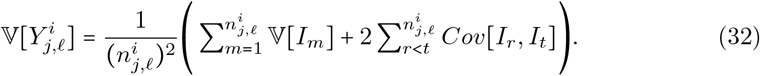

The variance of the indicator function *I_m_*

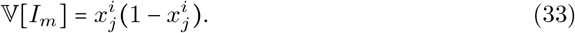

It remains to find the covariance

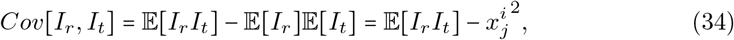

where, marginalizing on the underlying genotype *G* and assuming HWE, it follows that

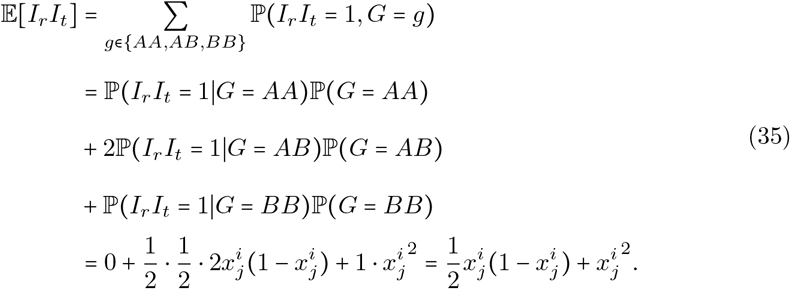

Considering that the sum over *r* < *t* in equation (32) is made over 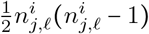 equal expectations, we can write

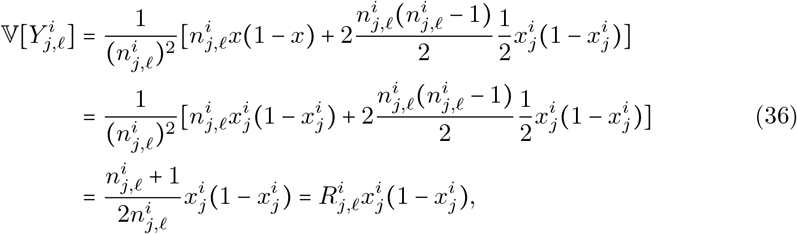

where for practical purposes we have defined, for each *ℓ*th individual, 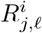 as the ratio

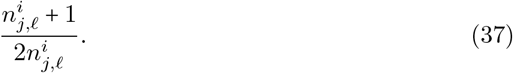

Consider at this point the approximation of the variance of the weighted “pseudo-individual”, having estimated frequency 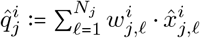.

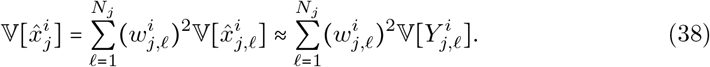

Our objective is to perform a Lagrange-constrained optimization w.r.t. the weights, being sure to find a minimum since eq (38), as function of the weghts, is convex. This is easily done since the Lagrange-parametrized function is

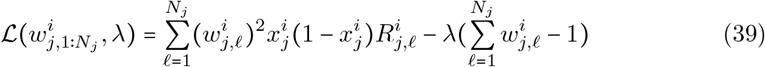

and it originates a linear system of equations of the form

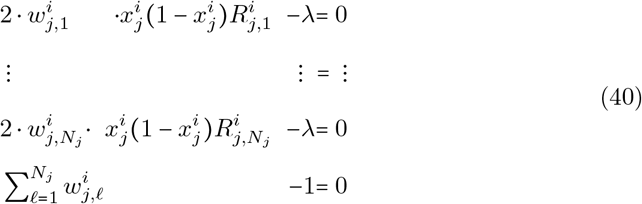

whose solution provides us with the minimum values of the weights as follows *∀ℓ ∊*{1,…,*N_j_*}:

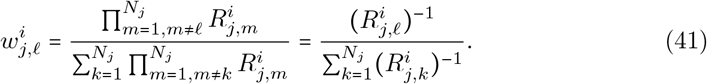

### Appendix 3 Error estimation and correction

Estimation of the type-specific errors follows the supplementary material of [19]. Assume having one observed sequenced individuals affected by base-transition errors. This individual has an associated 4x4 error matrix ***e***, such that the entry ***e***(*a, b*) is the probability of having sequenced allele *b* instead of allele *a*. Consider the tree ((T,R),O), in which the leaves are sequenced genomes affected by type-specific errors (T), an individual without errors, used as reference for the error correction (R), and an outgroup individual (O).

Assume that loci are independent and that the errors between pairs of alleles are independent given a base *o* in the outgroup and the error matrix ***e***. Then the likelihood of the base *t* in the observed individual can be decomposed as a product through the loci:

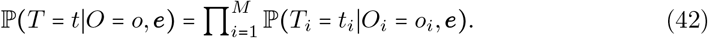

Marginalize any ith factor of the above equation over the true alleles before error *g_i_ ∊* {*A, C, G, T*} of the underlying true genotype:

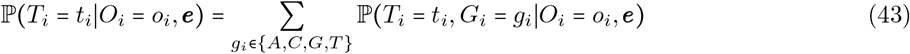

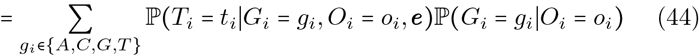

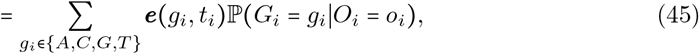

where the true genotype *g_i_* is independent of the error rates for each *i* = 1,…,*M*. One can approximate the probability of observing *g_i_* conditionally to o_i_ with the relative frequency of the base *g_i_* in the error-free individual *R*, for loci where the outgroup is *o_i_*, that is

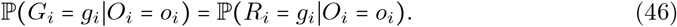

It is possible to perform a maximum likelihood estimation by numerical optimization to obtain an estimate of the error matrix. Note that every entry ***e**(g_i_,t_i_*) is the same over all loci.

The rationale behind the error correction is that the count of each base in the genomes T and R should be the same, otherwise an excess of counts in T is due to error. This approach to error estimation has been applied in [19] to study type-specific errors in ancient horses’ genomes.

Assume that the error matrix ***e**_ℓ_* has been estimated for every individual *ℓ* in each *j*th group. For a specific genome *ℓ* we have the following equation for each locus *i*

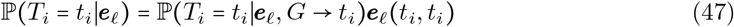

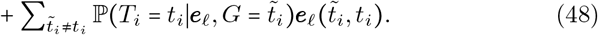

The same equation can be expressed in matrix form as follows:

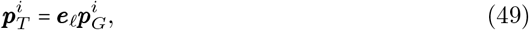

where 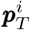 and 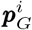 are the vectors of probabilities of observing alleles at locus *i*, respectively in the T and R genome. If the error matrix *e_ℓ_* is invertible, we can find the error corrected allele frequencies as

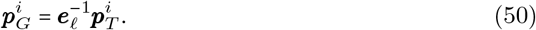

The correction performed in eq (50) makes the estimated allele frequencies unbiased. The unbiasedness allows the numerator of the D-statistic to have mean zero, and makes the D-statistic calculated with error-corrected frequencies convergent to a standard normal distribution (see Appendix 1). In fact, consider for a certain locus the di-allelic scenario with alleles A and B. Let *n* be the number of observed bases. The number of alleles A in absence of errors is

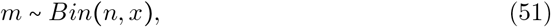

where *x* is the population frequency. Let *∊_A,B_* and *∊_B,A_* be the probabilities of having a transition from A to B and from B to A, respectively. Then the total number of observed A alleles is given by the sum of the two following variables:

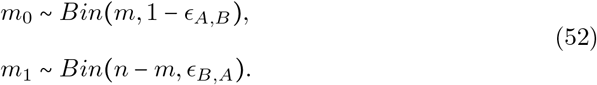

The expected population frequency is given by

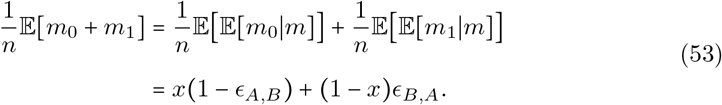

The error matrix and its inverse for the di-allelic case are expressed as follows:

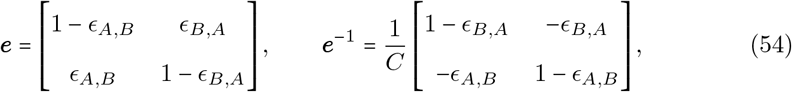

where *C* = (1 - *∊_A,B_*)(1 - *∊_B, A_*) - *∊_A,B_∊_B,A_* is the constant arising from the inversion of a 2 x 2 matrix.

The formula in eq (50) is rewritten as

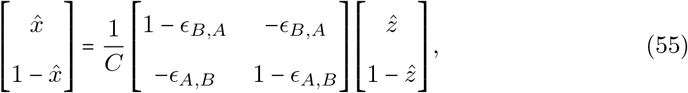

where 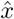 is the estimator of the error-corrected population frequency, while 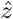 is the estimated population frequency prior to error correction:

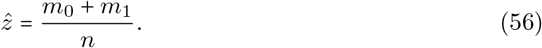

From eq (55) it is possible to deduce the following equality:

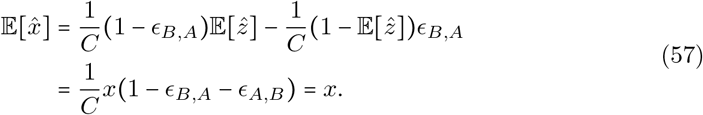

This proves that the error-corrected estimators of the allele frequencies are again unbiased, therefore calculating the D-statistic using error-corrected allele frequencies leaves the convergence results unchanged.

### Supplemental Data

The Supplemental Data contains two tables with numeric results related to a real data scenario, and three figures regarding the estimates of type-specific errors, the behaviour of the D-statistic and the correction for external introgression.

